# Colour preference in juvenile European lobster (*Homarus gammarus*)

**DOI:** 10.1101/2024.04.12.588567

**Authors:** Matt E. Bell, Nicholas Kuvaldin, Erik Romero Frontaura, Felix K.A. Kuebutornye, Paula Costa Domech, Alex H. L. Wan

**Affiliations:** Aquaculture and Nutrition Research Unit (ANRU), Carna Research Station, School of Natural 8 Sciences and Ryan Institute, University Galway, Carna, Co. Galway, Ireland, H91 V8Y1; Aquaculture and Nutrition Research Unit (ANRU), School of Natural Sciences and Ryan Institute, Annex building, University Galway, Carna, Co. Galway, Ireland, H91 TK33; University of South Bohemia in České Budějovice, Faculty of Fisheries and Protection of Waters, South Bohemian Research Center of Aquaculture and Biodiversity of Hydrocenoses, Institute of Aquaculture and Protection of Waters, České Budějovice, Czech Republic

## Abstract

Lobsters are high-value marine-fished species, and re-stocking efforts are essential for replenishing depleted wild populations and ensuring the longevity of these valuable marine species. The need for stock enhancement and hatcheries for aquaculture is vital to reduce the strain on wild populations. Therefore, environmental colours should be acknowledged when rearing lobsters in terms of behaviour, growth, movement, and survival. However, the behavioural effects the surrounding colour poses are virtually unknown. To better understand this behaviour, the movement of juvenile European lobsters (*Homarus gammarus*) was analysed through videos in two sub-trials. Lobsters were acclimated to custom-made alternating colour chambers, allowing for colour preferences based on individual movement. Sub-trial one alternated between black, red, blue, green, white, and yellow. The study identified that there was a significant preference for black (1 hour and 51 minutes) and red (1 hour and 33 minutes), indicating a potential underlying benefit for juvenile *H. gammarus* development. Despite this, little is known about the overall association of colour and lobster perception. A better understanding of environmental colour preference may guide the construction of aquaculture units to suit colour preference and enhance acclimation into newly released environments. This experiment aims to determine the preferred colours of stage four juvenile *H. gammarus* in a controlled laboratory setting and inspire further research to explore this unfamiliar field.

## 1. Introduction

Colours in the environment can significantly impact how animals interact with their environment and make important decisions for their survival (Twort and Stevens, 2023). In the aquatic environment, colour is a powerful language that organisms use to communicate and navigate their surroundings (Franklin et al., 2020). Previous research across various species has demonstrated how colour choices can influence behaviours such as mate selection (Detto et al., 2006), predator avoidance (Twort and Stevens, 2023), and habitat selection (Kawamura et al., 2020). For instance, studies on related crustaceans like fiddler crabs (*Uca mjoebergi*) and various stomatopod species have already revealed intriguing colour-related behaviours (Detto et al., 2006; Streets and Marshall, 2022), and in some species such as white leg (*Litopenaeus vannamei*), even the substrate colour has got an effect on their growth performance (Luchiari, Freire and Marques, 2012). The perception of colours exhibits variations across diverse species of aquatic animals, reflecting their unique visual sensitivities and adaptations to underwater environments, light conditions, structural complexity and their own behaviour, such as mating, feeding and activity patterns (diurnal and nocturnal) (De Buserolles et al., 2012).

The European lobster (*Homarus gammarus*, Linnaeus, 1758) is a commercially viable fishery worth over €40 million in 2021. However, the current wild European lobster (*Homarus gammarus*, Linnaeus, 1758) stocks are under pressure from many stressors, thus, reducing their fitness. Considerable efforts have been made to enhance stock in several European nations (Hinchcliffe et al., 2022). With a bottleneck rarely exceeding 20 % survival from the lobster planktonic larval phases (Addison and Bannister, 1994; Daniels et al., 2010; Ellis et al., 2015; Powell et al., 2017; Goncalves, 2021; Bell et al., 2023), stress impacts from environmental conditions, such as colour should be accounted for to reduce juvenile mortality within hatcheries. This is especially true during ecdysis, which is marked when larvae transition from stage II to III, where mortality rates peak (Addison and Banister, 1994).

In nature, colour is responsible for and intrinsically affects animal behaviour by attracting or deterring species settlement to specific habitats (Kelber and Osorio, 2010). Colours contrasting environmental backgrounds can also cue mating behaviour or signal a warning threat (Stevens and Ruxton, 2012). For example, the poison dart frog (*Dendrobatidae*) deters predators from feeding upon it by saliently evincing a contrasting blue, yellow, or red body (Wang and Shaffer, 2008). Concerning sexual selection, male blue crabs (*Callinectes sapidus*) prefer females with red claws over orange claws (Baldwin and Johnson, 2012). The colouration of *H. gammarus* is attributed to the environmental habitat setting but usually, it develops darker and bluer colouration (Tlusty et al., 2005). Therefore, understanding colour perception regarding natural selection, mating capability, and survivorship is a vital prerequisite for lobster autecology and should be considered a factor in conservation and restocking efforts. Lobster carapace colour is also economically and culturally associated, whereby uniform, red-coloured lobsters are more lucrative for open markets, providing the consumer luck, prosperity, and happiness (Konosu and Yamaguchi 1994).

The colour of cultivation tanks regulates stress factors, aggression, growth, and feeding alterations via hormonal and neural processes, resulting in excess metabolic output, and inhibited growth rates via catabolism (El-Sayed and El-Ghobashy, 2011). The colour of the cultivation tank affects light intensity and other light properties, such as reflectance and absorption (Lesmana et al., 2021). Extremely intense light for prolonged durations can induce stress signals and may lead to the death of cultivated species (Fitch and Lankford, 2013). Another study alternated between white and black backgrounds for *H. gammarus* juveniles, utilising Pollack (*Pollachius pollachius*) fish as a model for vision whilst monitoring carapace colourimetry (Mynott et al., 2018). In addition, darker colours (black) are preferred for aquaculture cultivation due to their positive photophobic response and orientation to reflective surfaces (Martin-Robichaud and Peterson, 1998). Alternatively, phototactic responses are induced in lighter colours (white).

Lobster juvenile rearing aspects merit further association of environmental colour preference and vision perception in cultivating lobsters, especially in aquacultural settings. This study aims to determine the most suitable environmental colour (red, green, blue, yellow, black, and white) for stage IV juvenile *H. gammarus* lobsters by measuring the timings of their movements from different colour chambers via video. Preferred colours may be replicated in future designs of aquaculture units to optimise development and simulate a more natural environment. This method will prompt physiological plasticity and acclimation of vulnerable juvenile lobster populations before release.

## 2. Materials and methods

### 2.1. Juvenile lobsters

Larvae were obtained from berried female lobsters sourced from a local commercial wholesaler. Stage I-III zoea was reared in 70 L hopper systems filled with pseudo-green seawater (mixture of *Nannochloropsis oculata, Dunaliella salina*, and *Isochrysis galbana*) and were fed daily with enriched artemia. Nearing to the end of stage III, lobster zoa were settled and reared in individual holding units and fed with particulate diets (C1/C2, Pacific Trading Aquaculture Ltd, Dublin, Ireland). When juveniles had fully settled to stage IV, 96 lobster juveniles (60 for sub-trial one, with an average length of 20.21

± 2.3 mm, carapace length of 9.27 ± 0.72 mm, width of 3.78 ± 0.35 mm and weight of 0.123 ± 0.02 g, and 36 sub-trial two, with an average length of 20.33 ± 1.24 mm, carapace length of 9.48 ± 0.55 mm, width of 3.42 ± 0.27 mm and weight of 0.129 ± 0.02 g) were randomly selected for the behavioural trial.

### 2.2. Design and production of experimental colour chamber

The colour preference trial was undertaken in colour chambers, which is comprised of six equally sized triangular sub chambers forming into a hexagon. An open area in the middle allowed the lobster to access the different colour sub-chambers. These sub-chambers consist of opaque side walls to minimise colour interference from other chambers besides the front entry point (Figure 1). The testing chambers were designed using Tinkercad software (Autocad, California, USA) and then printed using PLA+ (polylactic acid) on a Ultimaker S5 3D printers (Geldermalsen, The Netherlands) at the Make a Space unit, James Hardiman Library, University of Galway. For sub-trial one, the chamber comprises six different colour sub-chambers, i.e., red, blue, black, green, yellow, and white. While for sub-trial 2, it is made of 3 reds and 3 black sub-chambers. To ensure no colour bias, the arrangement of the sub-chambers was randomised.

**Figure 1.**
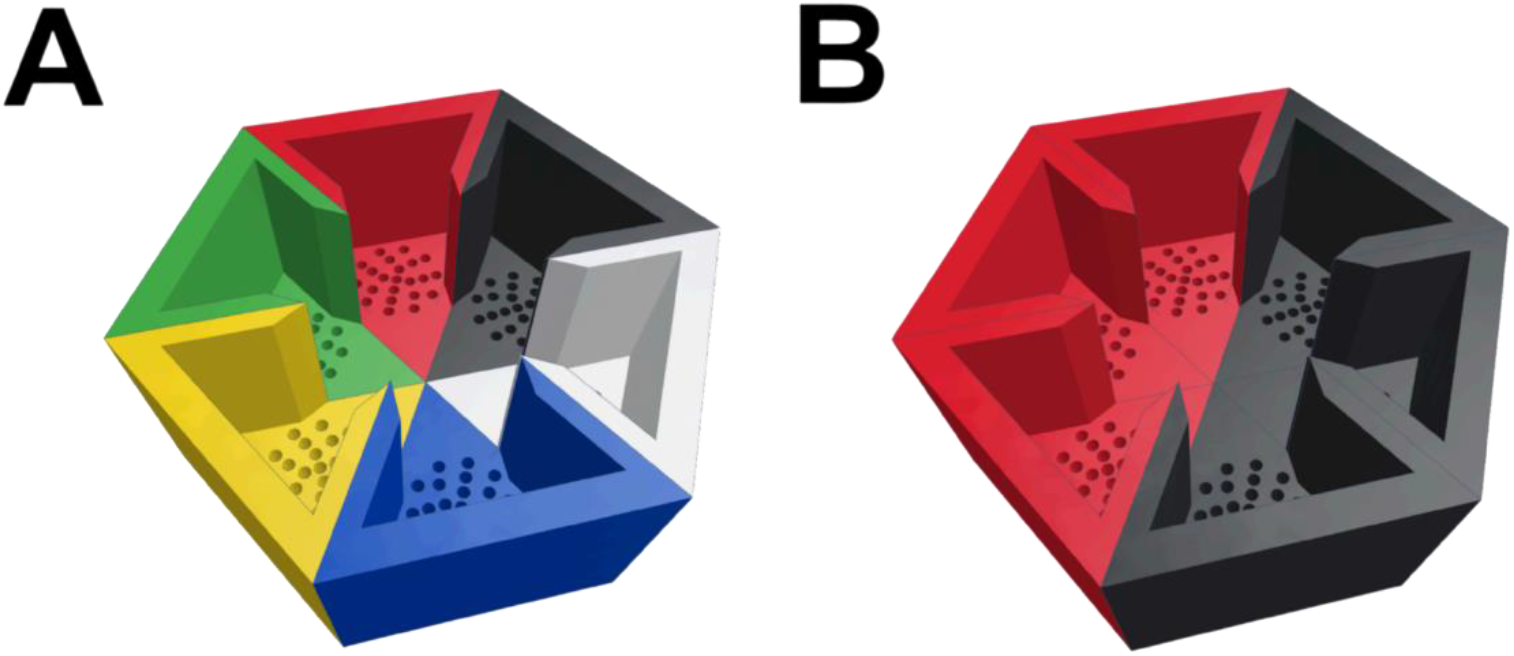
Illustration of the 3D printed colour chambers used in the colour behavioural trial. Each segment is printed individually to allow different colour permutation arrangements: A) Sub-chambers used for sub-trial one that is comprised of six different colours: red, blue, black, green, yellow, and white; B) Testing chambers used for sub-trial two comprised of red and black sub-chambers.

### 2.3. Trial design and setup

Each chamber was submerged in a tray of filtered seawater (35.40 ppt, 21.24 ± 1.01 °C), which was filled below the chamber walls to prevent escape. Fluorescent lighting units were suspended above the chambers to give a light lux intensity of 2731.37 ± 279.25 (Hobo MX temp/light system, ONSET, Bourne, MA, USA) and photon flux density was measured at 48.74 ± 2.95 µmol m^-2^ s^-1^ across the spectrum of 380-750 nm (LI180 spectrometer, LI-COR Biosciences UK, Cambridge, UK). An individual lobster was transferred into each chamber where it was acclimated in an obscured cylindrical tube in the centre of the chamber for 4 minutes before it was released, and the trial commenced. The colour preference trial was undertaken over 3 hours. To prevent bias preference behaviour (e.g., learned behaviour), an individual lobster undergoes the behaviour trial, it is not retested or reused in the second sub-trial.

### 2.4. Movement analysis

The lobsters’ movement during the trial was recorded using a Connect video camera (Logitech, Lausanne, Switzerland) that was mounted above the chamber and recorded onto a computer for later analysis. Colour preference for red, black, blue, yellow, green, and white colours was recorded when >50 % of the body was inside the sub-chamber. The colour and corresponding movement times were recorded, where the duration, incidental movements, and final colour settlement in colour chambers were calculated. The duration is how long the lobster spent within a colour chamber. The incidental movement is how many times the lobster entered the colour chamber. The final colour settlement was recorded based on the last colour chamber the lobsters settled in before the video concluded.

## 3. Results and Discussion

### 3.1. Colour preference

Extensive research has been focused on colour and its effects on animal behaviour, such as mating, foraging, survival, and habitat selection (Kelber and Osorio, 2010). Studies have primarily focused on crustacean colourimetry, particularly how carapace colour is modulated via artificially altering conditional parameters such as diet type, light irradiance, and temperature (Yeap et al., 2022). However, few papers have been dedicated to crustacean vision perception and how their behaviour can be influenced by the colour in their environment. The present study highlights for the first time the colour preference in the economically important European lobster (Macneil et al., 1997; van der Meeren, 2005). The findings will bridge the knowledge gap and assist in appropriately constructing rearing systems, considering colourimetry components for stock enhancement and optimum larval-rearing programmes for European lobster (Agnalt et al., 1999). It is essential to decipher whether the colour preference is an innate, inherent (genetic) characteristic in lobsters or if it is a memory technique, which is acquired through learning like in zebrafish (*Danio rerio*) (Roy et al., 2019).

#### 3.1.1. Colour effects on lobster behaviour

Red is considered an insensitive visual cue for marine species (Weiss et al., 2006). Lobsters observed in the spectral range of 520-525 nm green wavelength, which is shorter than the red wavelength of >600 nm (Kennedy and Bruno, 1961). However, red was preferred over green in this experiment (Figure 2). Red is rarely observed in the ocean due to absorption, which unexpectedly saw a relatively higher preference in sub-trial one (Figure 2; A, mean duration at 01 h:33 min) over other colours like blue (Figure 2; A, lowest mean duration at 00 h:09 min), which is extensively represented in oceans due to scattering effect (Stramski et al., 2004). A similar observation was made in shrimp (*L. vannamei*), where red and yellow were preferred over blue and green, resulting in increased feed intake and higher growth rate (Luchiari et al. 2012). Despite this knowledge of colour affecting the growth and appetite of shellfish, hatchery constructions and designs do not consider colour preference in the actual sense. The significant preference towards red may inherently correlate to the inner red pigment astaxanthin (Krawczyk and Britton, 2001). However, this would require further research, especially into the biochemical components of this observation.

**Figure 2.**
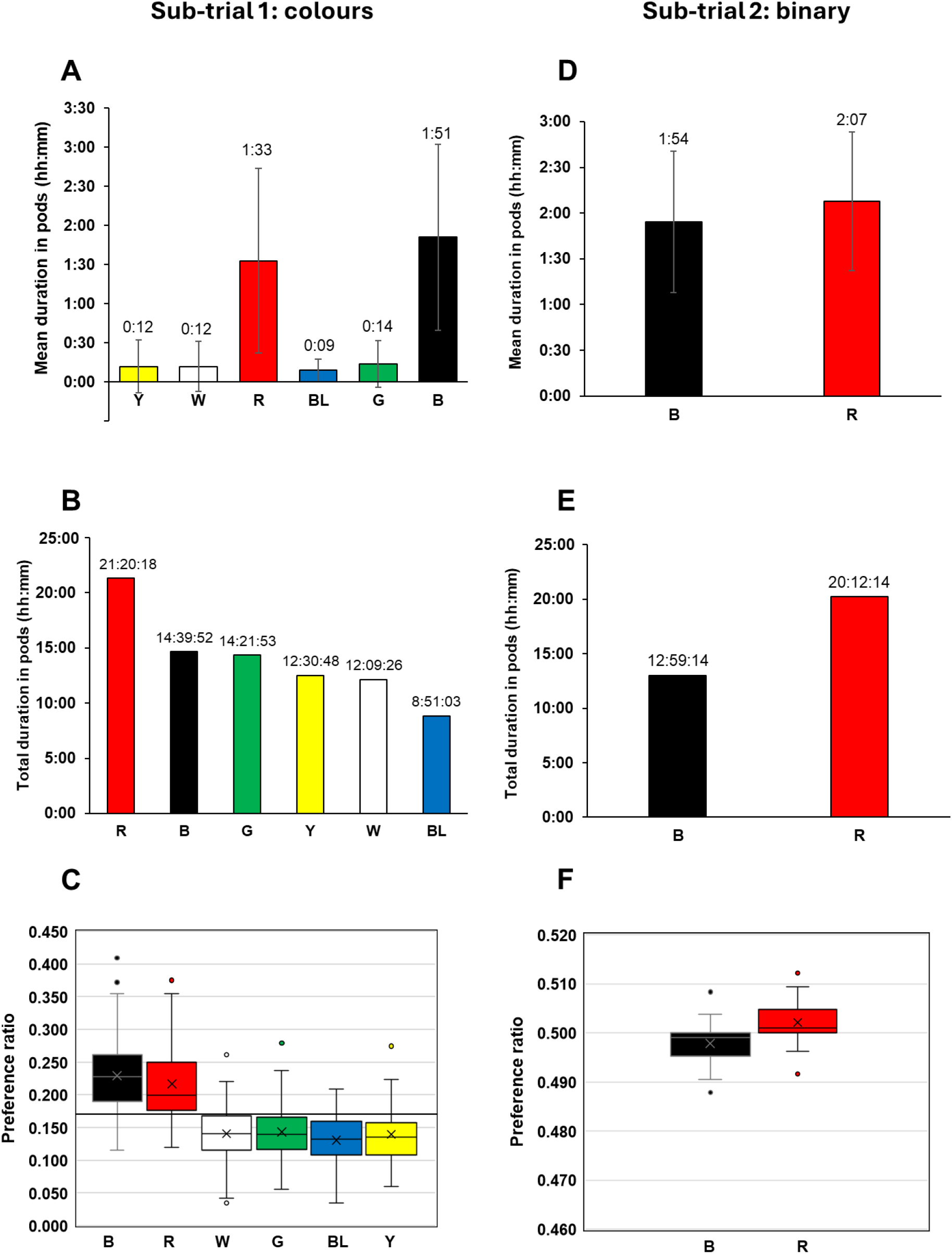
Colour preferences measured via various design parameters in both sub-trials: A) and D) Mean duration in hh:mm format; B) and E) Total duration in the colour chambers in hh:mm format; C) and F) Preference ratio, with a mean reference line (0.17) in figure C. Figure A, B, C, Sub-trial one, N=60; D, E, F, Sub-trial two N=36. B: black; BL: blue; G: green; R: red; W: white; Y: yellow.

It is also worth mentioning that red carapace colour is observed in Norway lobsters (*Nephrops norvegicus*) and American lobsters (*H. americanus*), both of which have a close phylogenetic relationship to *H. gammarus* (Pavičić et al., 2020). This elucidates that red colour preference might be an innate, evolutionary trait common to these lobster species. Another theory is that blue light poses an ‘awakening effect’, and red light promotes healing. This phenomenon is observed in humans, whereby exposure to blue light inhibits melatonin receptors, ultimately hindering circadian rhythm patterns of sleep (Wahl et al., 2019). Whereas red light is used as phototherapy to promote tissue recovery (Borges et al., 2014). The results might concur with the correlation between lobsters and their nocturnal and reposing lifestyle. The highest incidence of movement was for the black colour at 0.226 preference ratio in sub-trial one (Figure 2; C), which was expected due to the camouflage effect it produces on the lobster carapace. Darker backgrounds are preferred as potential ‘escape mechanisms’ from predation and threats. Whereas green and blue supposedly pose an inherent threat or deterrence from predators such as snapper (*Lutjanidae*), which exhibits blue, and octopus (*Octopus vulgaris*) capable of altering chromatophores to illuminate blue or green, depending on the situation (Veneza et al., 2023; Gilmore et al., 2016).

In comparison, the present results indicate inconsistency in colour preference between black and red, whereas in sub-trial one, black was preferred over red. However, in sub-trial two, red was most preferred. (Figure 2). Replicating this experiment will help to confirm the final colour preference, as the video analysis cut-off point does not account for a true representation of time. Both red and black were observed to have similar colour preferences in both sub-trials. In sub-trial one, the highest mean duration in pods was for black (01 h:51 min) followed by red (01 h:33 min), with a 14.19 % difference (Figure 2; A). However, when comparing the total duration time in the colour pods (Figure 2; B), red recorded the highest time with 21 h:20 min:18 sec compared to black (14 h:39 min:52 sec). Regarding preference ratio, there was a 4.06 % difference between black and red, with 0.23 and 0.22 preference ratios, respectively, in sub-trial two (Figure 2; E). On the other hand, in sub-trial two, red colour expressed higher values for all the parameters compared to black (Figure 2; D, E and F). However, the difference between preference ratios only accounted for 1 %, evincing high preference similarity.

Despite the results obtained in the present study, Lesmana’s (2021) study reported slightly different results for adult scalloped spiny lobsters (*Panulirus homarus*) in a similar methodological study. In lighting conditions, the highest preference was exhibited for black, followed by red. In contrast, the highest preference was expressed towards the red colour in nocturnal conditions, where movement into the red-coloured tank precipitated during nocturnal settings. However, the exact timings of the movement were not recorded here. Repeating this methodology under nocturnal conditions using *H. gammarus* will be necessary for variance or even alternating between light and night patterns for a genuine representation of the natural diurnal cycle. These darker conditions are preferred by lobsters, perhaps due to their nocturnal and solitary lifestyle, where they tend to hide in crevices to avoid predator detection in the wild (Lesmana et al., 2021).

Adult species of *H. gammarus* need consideration for ontogenetic factors in this type of experiment, as Lesmana et al. (2021) performed. Luchiari et al. (2012) discovered that Whiteleg shrimp (*Penaeus vannamei*) expressed the highest innate preference for red and yellow substrate after day three, where both colours exhibited enhanced feeding efficiency and specific growth rate, irrespective of surrounding colour. Moss et al. (1999) conducted a behavioural movement experiment of phyllosoma larvae under varying light intensities, measuring survival and growth. The researchers concluded that brightness affects aquaculture tank irradiance and food visibility. Optimum growth was observed under low light (0.001 µmol s^1^ m^2^) and dark (no light) conditions, compared to the highest light irradiance mode (10 µmol s^1^ m^2^). Future experiments should focus on brightness and irradiance measured separately as significant variables (Moss et al., 1999). In another experiment, Plichta et al. (2021) showed that Cherry shrimp (*Neocaridina davidi*) preferred darker backgrounds, regardless of carapace colouration. Suryanto et al. (2023) demonstrated that Louisiana crawfish (*Procambarus clarkii*) preferred red and deterred yellow colour, whereas Australian red claw crayfish (*Cherax quadricarinatus*) preferred blue and deterred yellow colour again. Kawamura et al., (2016), performed a colour preference experiment on giant freshwater prawn (*M. Rosenbergii*) based on chromaticity with three tones of grey. Sample shrimps were fed with beads of different colouration, with the highest preference observed in dark blue colour.

In a colour preference study, the next approach would be implementing various tones of colours, such dark red vs light red. Another suggestion is to incorporate unconventional secondary colours that are not normally present in the environment (e.g., orange, and purple), in tandem with primary colours for comparison (Dresp-Langley and Reeves 2018). Lobster zoea is phototactic, capable of utilising light to gauge and guide their movement towards food sources to increase intake and enhance development. Therefore, colour preference studies should be conducted to improve lobster larvae’s food intake and growth performance, focusing on lightness and darkness combined with feeding behaviour. Costa et al. (2023) suggested cultivating juveniles of the giant freshwater prawn (*Macrobrachium rosenbergii*) in blue surroundings and cultivating adult species in darker, lower-irradiance surroundings. This also highlights the need for a colour preference trial in adult H. gammarus.

#### 3.1.1. Colour impact on lobster physiology

Zheng et al., (2023) investigated growth rates, carapace colouration, mortality rates, and enzyme response in Australian freshwater crayfish (*C. quadricarinatus*). They observed darker backgrounds as preferable for *C. quadricarinatus* which is concurrent with this study’s findings on *H. gammarus*. Furthermore, Zheng et al. (2023) observed that amylase concentrations were highest in black and lowest in blue surroundings, likely due to crayfish’s foraging behaviour in colour-simulating their feeding habitat. This may also contribute to the observed growth advantages in crayfish cultured on a black background. The lowest reading for lipase was within blue and green surroundings, with superoxide dismutase activity at the lowest level within green but high in red surroundings. Additionally, catalase activity was highest in black and red and ceased in green surroundings (Zheng et al., 2023).

Future studies must investigate colour and the underlying mechanisms of enzymatic activities, especially in adult lobster populations. Resting periods, especially circadian rhythm and metabolism, must be supplemented in this investigation with performance correlation (Watson et al., 2022; Rodríguez-Viera et al., 2017). Overall, these studies disparately varied with the results obtained in this experiment. This is probably due to the utilisation of different species of crustaceans, which possess different spectral sensitivities to European lobsters. Similar enigmatic observations were made in stomatopods, evincing an extensive spectral range with poor selectivity for specific wavelengths (Streets et al., 2022). The study investigated the electrophysiological responses of photoreceptors, noting efficient selection for blue colours whilst not discerning red colour effectively after three months.

Chromaticity detection tests can also be applied to discriminate between colour hue or saturation components. This is to discern if colour is a visual factor or resembles a tone that is being perceived (Franklin et al., 2020). Future study perspectives must investigate lobsters’ colour preference and significance in animal welfare practises for restocking purposes and larval rearing. To better understand lobster ecology, aquaculture systems must mimic the natural environment of *H. gammarus*. Another approach is to conduct direct video analysis in the wild, as was demonstrated by Weiss et al., (2006), where lobster behaviour and movement were assessed on a benthic reef, utilising red light cameras, which are deemed insensitive to lobsters. This approach will provide direct observations of predation activity on vulnerable juveniles following release into the wild. Lobsters exhibit stress responses from high stocking densities during handling, transportation, and dissolved gas buildup. This may reduce meat quality as physiological parameters are hindered before the product even reaches the wholesaler. Certain colours may also induce physiological stress, making the lobster susceptible to pathological infection (i.e. vibriosis) by inhibiting immunological function (de Souza Valente and Wan, 2021).

## 4. Conclusion

Understanding the techniques required to implement and optimise juveniles for rearing and restocking purposes is still in early development. The present study highlights the colour preference in juvenile *H. gammarus* for the first time, providing substantial insight into how colour affects their movement and settlement preference. Red and black were significantly preferred over the latter in sub-trial one. Sub-trial two expressed a similar preference for black and red. This study recommends hosting lobsters in black, red or a mixture of both colours. However, scrutinising the preference between black and red will be required due to the high similarity in the binary layout of sub-trial two. This paper acts as a baseline for further research. Subsequent studies must focus on how colour correlates to behavioural traits in unison with feeding, welfare, stress, disease resistance, survival, and growth parameters.

## Acknowledgements

This study was conducted under the GLIOMACH seed fund project, which is funded by the College of Science and Engineering, University of Galway. The authors would like to acknowledge the technical support from Stephen McCusker, and Eileen Kennedy at Make a Space, Hardiman library, University of Galway for her assistance in the production of the 3D printed colour chambers.

## Author contributions

ERF, PCD, and AHLW contributed towards the experimental design. MEB, NK, ERF, PCD, FKAK, and AHLW undertook the study, data analysis and original writing. PCD and AHLW sorted the research funding. All authors made significant contributions toward this research study.

## Notes

### Competing Interest Statement

The authors have declared no competing interest.

